# Predicting driver distraction using a single channel ear EEG

**DOI:** 10.64898/2026.01.24.701469

**Authors:** Tzvetan Popov, Nan Li, Veronika Gambin, Kristina Keller, Samuel Wehrli, Stefan Lakämper

## Abstract

Cognitive distraction poses a major risk to driving safety, yet remains difficult to detect in real time because it often lacks overt behavioral markers. Neurophysiological measures such as electroencephalography (EEG) provide objective indices of cognitive load, but conventional EEG systems are impractical for real-world deployment due to their intrusiveness and computational demands. The present study examined whether cognitive distraction during naturalistic driving can be decoded using a single-channel, non-invasive in-ear EEG, and how its temporal dynamics compare with those obtained from full-cap scalp EEG and behavioral measures. Twenty-seven participants performed a dual-task paradigm in a highly immersive driving simulator, combining continuous vehicle control with low- and high-load arithmetic tasks presented on an in-vehicle display. Single-channel in-ear EEG, 24-channel scalp EEG, eye movements, and head rotation were recorded concurrently. Time-resolved multivariate pattern analysis was applied to decode working-memory load with millisecond precision across modalities. Cognitive distraction was reliably decoded from in-ear EEG, with detection latency and temporal generalization profiles closely matching those of full-cap scalp EEG. Although peak decoding performance was higher for scalp EEG, the timing and temporal stability of distraction-related neural signatures were largely overlapping between the two neural modalities. Eye velocity provided the earliest and most sensitive behavioral marker of distraction, while head rotation contributed complementary but weaker information. Scalp EEG topographies indicated that neural signals underlying decoding were closely linked to oculomotor and visuomotor processes engaged during task performance. These findings demonstrate that single-channel in-ear EEG provides a temporally precise and low-burden neural marker of cognitive distraction during driving. By prioritizing early detection and minimal hardware over maximal classification accuracy, the results identify a practical operating point for wearable EEG-based driver monitoring systems and support the feasibility of fast, off-the-shelf decoding approaches for real-world applications.

## Introduction

Driver distraction is a major contributor to road accidents and fatalities (Strayer et al., 2015). In particular, *cognitive* distraction- when a driver’s attention is diverted from the driving task to internal thoughts or secondary mental tasks- can severely impair driving performance even if the driver’s eyes remain on the road(Strayer et al., 2015). Unlike visual or manual distractions, cognitive distraction produces no obvious outward cues, making it difficult to detect and manage in real time. As modern vehicles integrate more infotainment and communication systems, there is growing concern that cognitive distraction will increasingly compromise traffic safety unless objective monitoring methods are developed (Prabhakar et al., 2018).

Researchers have turned to neurophysiological and behavioral measurements to objectively assess driver cognitive state. Electroencephalography (EEG) in particular has long been recognized as a powerful tool for indexing mental workload and attentional states during driving (Sonnleitner et al., 2014b; Wang et al., 2018; Peng et al., 2022; Li et al., 2023; Ronca et al., 2024). Characteristic changes in brain oscillations accompany distracted driving - for example, elevations in frontal theta and concomitant reductions in alpha power have been observed under cognitive load(Li et al., 2023). Such EEG-derived metrics can distinguish distracted versus attentive states, enabling the concept of a “distraction index” derived from brain activity(Ronca et al., 2024). Peripheral physiological signals likewise reflect cognitive distraction: pupil diameter is a well-known index of mental effort, dilating reliably with increasing working memory load(Prabhakar et al., 2018). Eye-tracking measures have shown that cognitive load often narrows the driver’s visual attention (e.g. longer gazes toward a task screen and fewer mirror checks), while head movements may indicate compensatory scanning or orienting when attention lapses(Reimer et al., 2012; Nilsson et al., 2020; Broadbent et al., 2023). Indeed, high workload dual-tasks can produce prolonged off-road glances and increased corrective head rotations as drivers struggle to divide attention(Reyes and Lee, 2008; Mikula et al., 2020). These converging lines of evidence have spurred interest in *multimodal* monitoring approaches, combining neural and peripheral measures to improve sensitivity. Notably, a recent review concludes that integrating EEG with other physiological signals is a promising avenue for enhanced driver distraction detection(Li et al., 2023). Furthermore, distraction-related changes in these signals can manifest rapidly- within as little as one second of increased cognitive load(Prabhakar et al., 2018)- underscoring the importance of time-sensitive analysis methods for capturing transient lapses of attention.

For real-world usability and practical applicability, there is a clear need for EEG measures that are readily accepted by drivers and impose minimal burden on natural behavior. Ideally, such measures should rely on unobtrusive, everyday form factors- such as ear-bud-based or ear-EEG systems- which resemble common contemporary consumer devices and are therefore more likely to be tolerated in applied settings. Equally important, the extracted neural information should require minimal preprocessing and still provide robust markers of cognitive distraction that are comparable in sensitivity and interpretability to established behavioral or physiological indicators. However, most existing studies of driver distraction have relied on unimodal measures and simplified experimental paradigms with limited ecological validity. Many laboratory investigations employ abstract or isolated secondary tasks (e.g., mental arithmetic or vigilance tasks) under highly controlled conditions, often using static driving simulators or even omitting an active driving task altogether. While such approaches are informative for isolating specific cognitive processes, they fail to capture the full complexity and dynamic temporal evolution of distraction as it unfolds in real-world driving. Although hybrid approaches combining EEG with eye tracking and other behavioral sensors have frequently been advocated, relatively few studies have explicitly linked neural activity to naturalistic gaze behavior and head movements during driving. As a result, our understanding of how cognitive distraction concurrently affects neural, ocular, and motor systems in real time remains limited. This gap in multimodal, time-resolved research constrains both theoretical progress and translational application. Consequently, there is a pressing need for studies that investigate driver distraction in immersive, behaviorally rich environments, using analytical approaches capable of decoding evolving cognitive states from multiple synchronized data streams. In particular, demonstrating that low-intrusion EEG measures can reliably index distraction alongside natural eye and head movement behavior would represent a critical step toward scalable, real-world neuroadaptive driving systems.

Here, these challenges are addressed by leveraging time-resolved multivariate pattern analysis (MVPA) based on linear discriminant analysis (LDA) applied to multimodal biosignals in a highly immersive simulated driving environment. Time-resolved MVPA provides a principled means of quantifying moment-by-moment neural and physiological patterns that discriminate between high and low cognitive load states. In the present study, single-channel in-ear EEG, 24-channel scalp EEG, together with eye movement measures (gaze velocity) and head rotation data were recorded, while participants performed a dual task in a realistic full-scale driving simulator. The secondary task required participants to solve difficult versus easy mental arithmetic problems presented on the central dashboard display while maintaining control of the vehicle.This dual-task paradigm required drivers to direct visual and cognitive attention toward the middle dashboard in order to perceive, evaluate, and respond to the arithmetic problems, thereby inducing graded levels of cognitive distraction and working-memory load concurrent with driving. Importantly, this manipulation preserves ecological validity, as it closely resembles common in-vehicle interactions such as engaging with navigation or infotainment systems.

By uniquely integrating multiple physiological modalities within a time-sensitive decoding framework and an immersive driving context, this approach captures the dynamic interplay between brain activity, gaze behavior, and head movements during cognitive distraction. In the following, this framework is used to demonstrate how objective neurophysiological and behavioral markers of driver distraction can be identified, and its implications for neuroergonomic monitoring of cognitive states in real-world driving scenarios are discussed.

## Methods

### Participants

Twenty-seven individuals took part in the study. Participants were recruited via the local university as well as through public community advertisements. The resulting sample included 18 women and 9 men, ranging in age from 22 to 50 years (mean age = 28.6 years). All participants provided written informed consent prior to participation, in accordance with the Declaration of Helsinki. Ethical approval for the study was obtained from the local institutional review board of the host institution (IRB no. 25.05.15). Participants received financial compensation for their time.

The present study was not preregistered. The achievable sample size was determined by practical considerations, including time and budgetary constraints. As discussed by (Lakens, 2022), such constraints should be explicitly acknowledged when an a priori power analysis cannot be fully implemented, and this limitation is therefore transparently reported here. While the present analyses partially rely on data reported in earlier work(Popov et al., 2025), the research questions, analytical approach, and interpretation presented here are distinct and non-overlapping.

### EEG and Eye Tracking

Visual stimuli were presented on a Fujitsu CELSIUS H-Series monitor (1920 × 1080 resolution). Participants were seated in the driver’s position behind the steering wheel of a BMW i3 vehicle mock-up. Stimulus presentation was controlled using PsychoPy (Peirce et al., 2019) and displayed full-screen against a black background.

In-ear EEG data were recorded from all 27 participants using a device developed by IDUN Technologies (https://iduntechnologies.com/). The in-ear EEG constitutes the primary data source of the present study and forms the basis of all main analyses reported here. In addition, multichannel scalp EEG was recorded using a wireless 24-channel mBrainTrain SMARTING system equipped with semi-dry saline-based Ag/AgCl electrodes mounted in an elastic cap and arranged according to the international 10-20 system. Signals were referenced online to FCz, with AFz serving as the ground electrode. Electrode impedances were maintained below 10 kΩ. Data were sampled at 250 Hz and re-referenced offline to the common average. The cap EEG served a secondary role, providing reference information on the temporal evolution and scalp topography of task-related neural activity to support interpretation of the single-channel in-ear EEG signals.

Eye-movement data were collected using Tobii Pro Glasses 3. The system recorded binocular gaze positions and pupil diameter, as well as head-motion signals derived from integrated gyroscope and accelerometer sensors. All eye-tracking data were streamed in real time via the Lab Streaming Layer (LSL) protocol. Event markers corresponding to stimulus onset were transmitted through a dedicated LSL stream to ensure precise temporal synchronization across EEG, eye-tracking, and SILAB (see below) data streams.

### Driving simulator setup

The driving experiment was conducted using the VICTOR (Vehicle for Interdisciplinary Clinical and Translational Research) simulator, which is based on a fully integrated BMW i3 vehicle platform. The original suspension system was replaced with custom-built mounts supporting electrically driven spindle actuators, enabling controlled motion feedback primarily related to road surface irregularities, acceleration, and braking. Several custom-manufactured components were incorporated to support this motion system. Standard mirrors and dashboard elements were either concealed or replaced by LCD displays (Beetronics 10HD7 and 13HD7). A 40-inch display (IIYAMA X4071UHSU-B1) mounted in the rear section of the vehicle simulated the rear-view mirror, while an additional touch-sensitive display (Asus Zen Touch MB16AMT) was integrated into the center console to present task-relevant information and record responses. Steering, indicator controls, and pedal systems were adapted to interface directly with the simulation software. Vehicle dynamics and the virtual driving environment were generated using SILAB (version 7.0; WIVW, Würzburg, Germany). Visual output was rendered both on the in-vehicle displays and via five laser-phosphor DLP projectors (Barco F-80_Q7), which projected onto a fixed 275° curved screen. Image warping and blending were handled using VIOSO AnyBlend software. The simulation was rendered and logged at 60 Hz. The system operated on a network of ten identical computers controlled centrally via SILAB. Environmental conditions in the simulator room were maintained at a stable temperature, and the vehicle cabin was continuously ventilated with fresh air. Most technical hardware was concealed from view, resulting in a visually minimal setup resembling a stationary vehicle in a garage environment, thereby reducing potential discomfort or apprehension in participants.

### Driving scenario and behavioral measures

Participants completed a customized nighttime highway-driving scenario designed to induce low sensory stimulation over an extended period. The scenario was composed of ten sequential modules derived from an established vigilance-driving paradigm used in standardized driver fitness assessments. Across approximately 45 minutes, participants followed a lead vehicle on a single-lane roadway without overtaking. The lead vehicle varied its speed between 80 and 90 km/h, while the nominal speed limit remained constant at 100 km/h. Oncoming traffic appeared intermittently at variable intervals. To minimize visual distraction and light emission, the vehicle’s speedometer was disabled. Motion cues were restricted to surface changes and speed variations; no additional motion feedback was provided for lane departures or collision events. Driving performance for each scenario segment was automatically evaluated using SAFE (Standardized Application for Fitness to Drive Evaluations), which interfaces directly with SILAB. Continuous driving metrics-including vehicle speed, headway distance, and standard deviation of lateral position- were recorded within SILAB and streamed via Lab Streaming Layer (LSL) for synchronization with EEG and eye-tracking data. In addition, discrete driving events such as lane departures, near-collisions, and collisions were automatically logged by the SAFE system.

### Experimental procedure

Participants were first familiarized with the simulator during a brief 10-minute practice drive to ensure comfortable handling of the vehicle controls and to screen for simulator sickness. No adverse symptoms were reported during this phase. The experimental protocol was then explained in detail, after which participants provided written informed consent. Subsequently, EEG equipment was prepared and signal quality verified, followed by fitting and calibration of the eye-tracking system (Tobii Pro Glasses 3). Participants were then seated in the simulator, and a final synchronization check across EEG, eye tracking, and driving data streams was conducted using Lab Streaming Layer. The experimental session consisted of the monotonous nighttime driving task, during which arithmetic problems were presented on a dashboard-mounted display under conditions of low and high working-memory load. These stimuli were shown in randomized order while participants continued to drive. Neural, ocular, head-motion, and driving-performance data were recorded continuously throughout the session. After completion of the approximately 45-minute drive, participants were debriefed, all equipment was removed, and monetary compensation was provided.

### Data Processing and Analysis

All analyses were conducted in MATLAB using the FieldTrip toolbox(Oostenveld et al., 2011). Data streams from in-ear EEG, scalp EEG, eye movements, head kinematics, and driving-related signals were temporally synchronized via Lab Streaming Layer and segmented into epochs time-locked to stimulus onset (-1 to 5 s).

#### Preprocessing

In-ear EEG data were demeaned and detrended, followed by visual inspection for artifact rejection. Trials with excessive noise were removed prior to further analysis. For event-related analyses, signals were baseline-corrected using the -1 to 0 s pre-stimulus interval. Eye-movement measures were transformed into gaze velocity, and head-rotation signals were extracted from inertial sensors. Missing samples in peripheral signals were handled conservatively, with short gaps interpolated and longer gaps retained as missing values prior to aggregation. All peripheral signals were epoched and baseline-corrected using the same temporal window as the EEG data to ensure comparability across modalities.

Scalp EEG data were preprocessed separately and used to characterize the temporal evolution and spatial distribution of task-related neural responses. These data served a secondary, interpretative role and were not the primary focus of decoding analyses.

### Event-related analyses

Event-related potentials (ERPs) were computed separately for low (easy) and high (hard) working-memory load conditions. For visualization and statistical summaries, ERPs were averaged across trials within participants and subsequently across participants. Scalp EEG topographies were used to illustrate condition-specific differences in amplitude and time course across the sensor array.

### Time-resolved multivariate pattern analysis

Time-resolved multivariate pattern analysis (MVPA) was applied to assess when neural and peripheral signals discriminated between high and low cognitive load. MVPA was implemented using linear discriminant analysis (LDA) classifiers within FieldTrip’s MVPA framework. Classifiers were trained and tested on time-resolved amplitude features only. Decoding was performed in a sliding-window manner across the entire epoch to obtain moment-by-moment estimates of classification performance. Performance was quantified using area under the receiver operating characteristic curve (AUC), with chance level defined as 0.5. To improve signal-to-noise ratio and target population-level distraction patterns, classifiers were trained on data pooled across all participants, rather than within individual participants. Cross-validation was implemented using k-fold (10 folds) procedures with multiple repetitions (5 repetitions) to ensure robust performance estimates.

### Temporal generalization analysis

To examine the temporal stability of distraction-related patterns, temporal generalization matrices were computed by training classifiers at one time point and testing them at all other time points. This analysis revealed whether neural and behavioral signatures generalized across time or were transient and time-specific.

### Multimodal comparison

Decoding analyses were conducted separately for each modality (eye velocity, head rotation, in-ear EEG, and scalp EEG). Resulting AUC time courses and temporal generalization matrices were compared to assess the relative sensitivity and temporal dynamics of each signal source. This multimodal approach enabled direct comparison of neural and peripheral markers of cognitive distraction within a unified analytical framework.

### Statistical Analysis

To control for multiple comparisons, statistical testing was performed using a cluster-based permutation approach (Maris and Oostenveld, 2007). This method accounts for spatial, temporal, and frequency correlations by identifying clusters of adjacent significant data points. A two-tailed alpha threshold of 0.05 and 1000 permutations were used to assess statistical significance.

### Data and Code Availability

All data and the code necessary to reproduce the present results will be made available upon publication.

## Results

An overview of the task from the driver’s point of view is shown in Figure 1. During the driving task, participants followed a lead vehicle on a single-lane highway and were instructed to keep their gaze on the road ahead, as indicated by the eye-tracking cursor (red circle). Periods in which gaze remained directed toward the roadway were treated as the baseline condition in the present analyses. At pseudo-random intervals, an arithmetic stimulus was presented on a centrally positioned display located to the right of the driver. Stimuli were designed to impose either low or high working-memory (WM) load, hereafter referred to as *easy* and *hard* conditions, respectively. Presentation of the stimulus reliably elicited a gaze shift away from the roadway toward the display, illustrated by the displacement of the eye-tracking cursor in Figure 1. Participants indicated whether the displayed solution was correct by pressing a response button once for correct solutions or twice for incorrect solutions (Figure 1A, right). Low-load trials involved simple arithmetic (e.g., *5 + 3 = 9*), whereas high-load trials required more complex operations (e.g., *8 + 6 × 6 = 44*). Across the approximately 45-minute driving session, a total of 200 stimuli were presented (100 per condition) in randomized order. Working-memory load had a clear impact on behavioral performance. Responses were faster in the low-load condition (M = 3.2 s, SD = 1.1) than in the high-load condition (M = 4.6 s, SD = 1.6), a difference that was statistically significant, *t*(26) = −8.9, *p* = 1.9 × 10⁻⁹, 95% CI [−1.67, −1.05]. Accuracy was also higher for low-load trials (M = 98.1%, SD = 1.95) compared with high-load trials (M = 95.5%, SD = 3.4), *t*(26) = 4.7, *p* = 7.32 × 10⁻⁵, 95% CI [1.47, 3.77].

**Figure 1:**
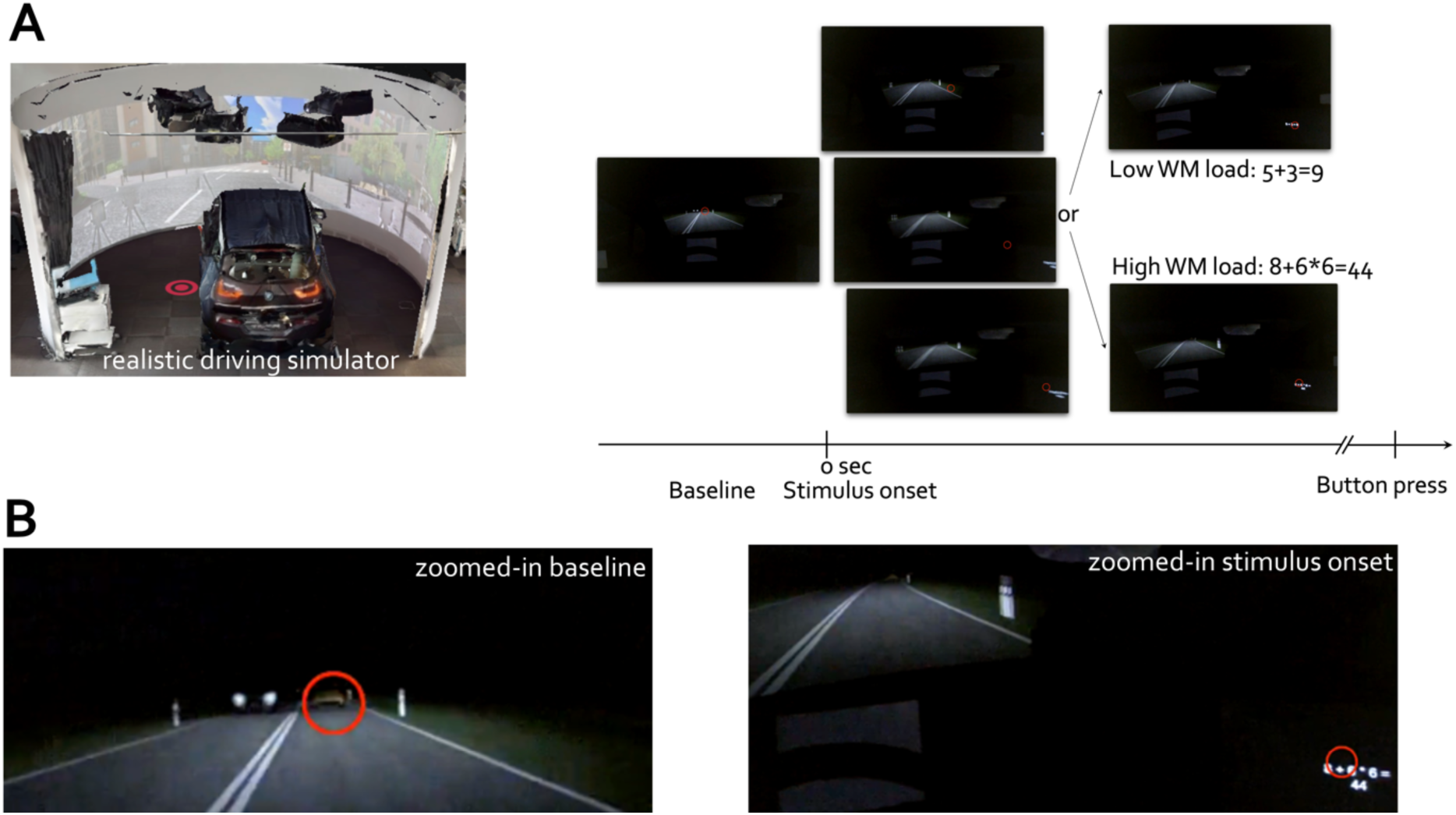
Dual-task paradigm combining simulated driving with an arithmetic working memory (WM) task, illustrating performance decrements under high WM load. **A-** Illustration of the realistic driving simulator. Schematic timeline of a trial, showing an initial baseline period followed by stimulus onset and presentation of an arithmetic problem of either low (e.g., 5 + 3 = 9) or high WM load (e.g., 8 + 6 × 6 = 44). Participants had to indicate via button press whether the presented solution was correct (one space key press) or incorrect (double space key press) while navigating a simulated driving environment. Red circles indicate the participant’s gaze position (point of view, 1dva) in the driving scenes. **B-** zoomed-in views of the gaze positions for the baseline (left) and stimulus (right) periods, highlighting the precise visual focus of participants during each task phase.

### Temporal evolutions of distraction-related responses

Figure 2A-D illustrates the stimulus-locked time courses of eye movement, head movement, and neural signals under low and high working-memory (WM) load.

**Figure 2:**
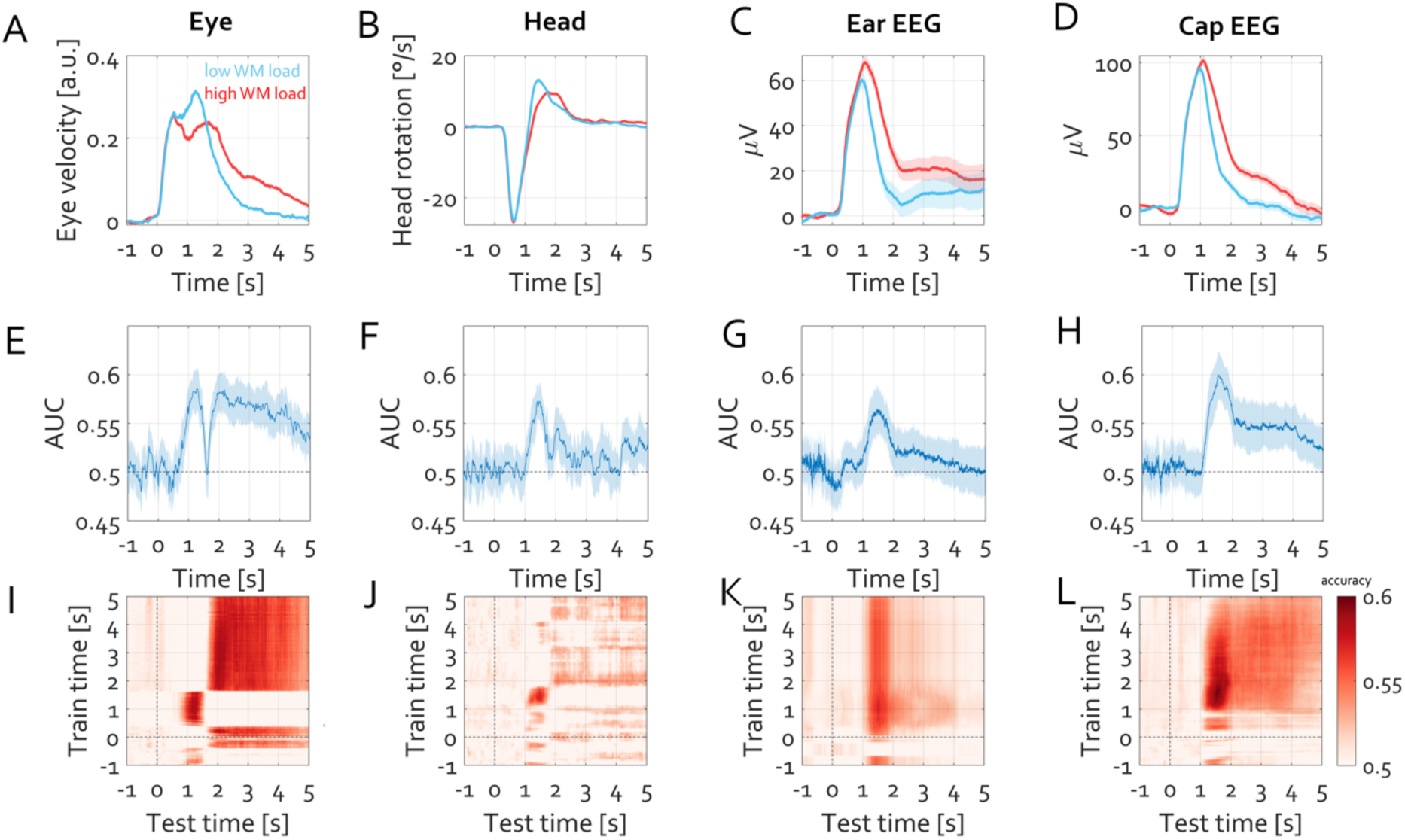
Multimodal neural and behavioral signatures of cognitive load and their time-resolved discriminability. **A-D:** Stimulus-locked time series for low (easy; blue) and high (hard; red) working-memory (WM) load conditions. **A-** Eye-movement responses quantified as gaze velocity show a rapid increase following stimulus onset, with a more sustained elevation under high WM load. **B-** Head rotation velocity (yaw axis) exhibits an initial rotation toward the dashboard stimulus followed by a return toward the road; this return is attenuated and delayed under high WM load, indicating prolonged head orientation toward the secondary task. **C-** In-ear EEG responses display a prominent stimulus-evoked potential followed by sustained activity, which is stronger for high WM load. **D-** Scalp EEG event-related potentials show a comparable temporal profile, with enhanced sustained amplitudes under high compared to low WM load. Shaded regions indicate ±1 SEM across participants. **E-H:** Time-resolved decoding performance (area under the curve; AUC) for discriminating high versus low WM load using linear discriminant analysis (LDA) applied separately to each modality. **E-** Eye velocity, **F-** head rotation, **G-** in-ear EEG, and **H-** scalp EEG all show above-chance classification performance following stimulus onset, with modality-specific temporal profiles. Shaded regions denote variability across cross-validation folds.**I-L:** Temporal generalization matrices illustrating classification accuracy as a function of training time (y-axis) and testing time (x-axis) for each modality. **I-** Eye velocity, **J-** head rotation, **K-** in-ear EEG, and **L-** scalp EEG. Warmer colors indicate higher classification accuracy. Sustained off-diagonal generalization suggests temporally stable representations of cognitive load. Vertical and horizontal dashed lines mark stimulus onset.

Eye movement dynamics (Figure 2A) showed a rapid increase in gaze velocity following stimulus onset, reflecting the saccadic shift away from the roadway toward the dashboard display. This initial peak occurred within 400ms after stimulus onset and was followed by a sustained elevation in gaze velocity. Notably, gaze velocity remained higher and decayed more slowly in the high WM load condition compared to the low WM load condition, indicating prolonged visual engagement with the task-relevant stimulus under increased cognitive demand. Head movement responses (Figure 2B) revealed a stereotypical pattern of head rotation toward the stimulus location, with angular velocity peaking approximately 800 ms after stimulus onset. This was followed by a compensatory rotation back toward the forward driving direction around 1.5 s post-stimulus. Under high WM load, the return rotation was attenuated and temporally delayed relative to the low-load condition, suggesting sustained head orientation toward the dashboard display and reduced reorientation toward the road. In-ear EEG responses (Figure 2C) exhibited a pronounced event-related deflection following stimulus onset, peaking within the first 1.5 s and followed by a sustained neural response lasting several seconds. While early components were comparable across conditions, the sustained activity was markedly stronger for the high WM load condition, indicating prolonged engagement during cognitively demanding trials. Scalp EEG responses (Figure 2D) showed a similar temporal profile, with a large stimulus-evoked potential followed by sustained activity extending several seconds post-stimulus (averaged across all recording electrodes). As observed in the in-ear recordings, sustained EEG amplitudes were greater under high WM load than low WM load, suggesting that increased cognitive demand modulated prolonged processing beyond the initial sensory response.

Together, these results demonstrate converging effects of working-memory load across oculomotor, head kinematics, and neural signals, with high cognitive demand consistently associated with prolonged gaze shifts, delayed head reorientation, and sustained neural activity following stimulus onset.

### Decoding of cognitive load across modalities

Time-resolved multivariate decoding was used to assess when neural and peripheral signals discriminated between low and high working-memory (WM) load. Decoding performance is reported as area under the receiver operating characteristic curve (AUC), with chance level at 0.5. Across modalities, decoding accuracy increased reliably following stimulus onset (Figure 2E-H). Eye-movement-based decoding (Figure 2E) showed the earliest rise above chance, beginning shortly after stimulus onset and reaching peak performance around 1.2s post distractor onset. Decoding performance remained elevated for several seconds, indicating that gaze dynamics carried sustained information about cognitive load.

Head-rotation-based decoding (Figure 2F) also exceeded chance level, though with a slightly delayed onset and lower peak performance compared to eye movements. Classification accuracy remained modest but consistently above chance during the post-stimulus interval, suggesting that head-orienting behavior provided complementary, though weaker, information about distraction level.

In-ear EEG-based decoding (Figure 2G) revealed a clear increase in classification performance following stimulus onset, peaking around 1-2 s and gradually declining thereafter. While overall AUC values were lower than those obtained from scalp EEG, decoding performance was robustly above chance for an extended period, demonstrating that single-channel in-ear EEG contained discriminative information about WM load.

Scalp EEG-based decoding (Figure 2H) yielded the strongest and most sustained classification performance among all modalities. AUC increased sharply after stimulus onset, reached a pronounced peak around 1-2 s, and remained elevated for several seconds, indicating stable neural signatures of cognitive load.

Temporal generalization analyses further characterized the stability of these effects (Figure 2I–L). Eye velocity exhibited robust off-diagonal generalization across an extended post-stimulus interval, indicating that the discriminative gaze patterns associated with cognitive load were temporally sustained rather than confined to brief, time-specific events. Once established following stimulus onset, these gaze-related patterns generalized broadly across time, consistent with prolonged visual engagement with the secondary task. Head-rotation decoding showed more limited temporal generalization, with performance strongest along the diagonal and weaker off-diagonal transfer, suggesting a more transient and time-specific contribution of head-orienting behavior. In contrast, both in-ear EEG (Figure 2K) and scalp EEG (Figure 2L) exhibited pronounced off-diagonal generalization, reflecting temporally stable neural representations of cognitive load that generalized across multiple post-stimulus intervals. Together, these results indicate that eye velocity and neural signals share a high degree of temporal stability, whereas head-rotation signals reflect more temporally constrained dynamics.

Together, these results demonstrate that cognitive load during driving can be decoded from multiple signal sources, with eye movements providing early and sensitive markers, neural signals supporting more temporally stable representations, and in-ear EEG offering a viable, low-intrusion neural measure whose decoding performance approaches that of conventional scalp EEG.

### Latency comparison across modalities and topographic characterization of neural decoding signals

Figure 3 summarizes the relative timing of distraction-related information across behavioral and neural modalities and illustrates the scalp distribution of the corresponding neural signals. Figure 3A shows the time-resolved decoding performance for eye velocity, head rotation, in-ear EEG, and scalp EEG, plotted together to facilitate direct latency comparison.

**Figure 3:**
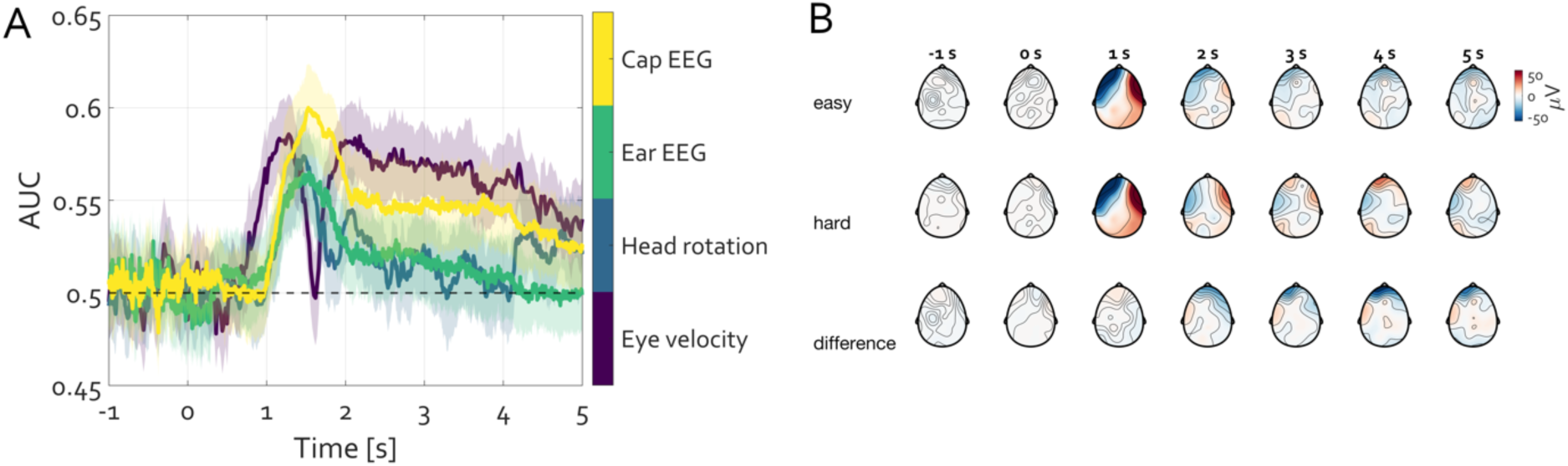
Latency comparison of distraction-related decoding across modalities and scalp topography of neural effects. **A-** Time-resolved decoding accuracy (AUC) for discriminating high versus low working-memory load based on eye velocity, head rotation, in-ear EEG, and scalp EEG. Eye velocity shows the earliest rise above chance, followed by head rotation. In-ear EEG and scalp EEG exhibit closely matched onset latency and temporal profiles, indicating comparable timing of neural information despite the difference in sensor count. Shaded areas denote variability across cross-validation folds; dashed line indicates chance level (AUC = 0.5). **B-** Scalp topographies at 1-s intervals relative to stimulus onset for the easy condition (low WM load, top row), hard condition (high WM load, middle row), and their difference (bottom row). Topographies reveal sustained posterior and lateral distributions of task-related activity, suggesting that neural signals contributing to decoding are closely linked to oculomotor and visuomotor processes engaged during task performance.

Decoding based on eye velocity rose earliest above chance, beginning shortly after stimulus onset and peaking within the first second. Head-rotation decoding followed a similar but delayed trajectory, with lower peak accuracy and a more gradual temporal profile. Both behavioral measures thus provided early indicators of task engagement, consistent with rapid oculomotor and orienting responses toward the secondary task.

In-ear EEG decoding exhibited a later onset than eye and head movement measures, with AUC increasing reliably above chance approximately 1-1.5 s after stimulus onset. Notably, the latency and overall temporal profile of in-ear EEG decoding closely resembled that of scalp EEG. Although peak decoding accuracy was lower for in-ear EEG than for the full cap EEG, the rise time, peak latency, and subsequent gradual decline were largely overlapping across the two neural modalities. This temporal correspondence indicates that single-channel in-ear EEG captures task-related neural dynamics with a timing comparable to that obtained from multichannel scalp recordings.

To further characterize the neural sources contributing to decoding, Figure 3B presents scalp topographies of stimulus-locked activity at 1-s intervals for the easy and hard conditions, as well as their difference. Across post-stimulus time points, both conditions showed pronounced spatial patterns over frontal and lateral scalp regions consistent with the timing of maximal change in eye velocity and subsequent head rotation. The difference maps revealed sustained modulations with a distribution also over lateral and frontal electrodes.

Together, these results demonstrate that single-channel in-ear EEG provides a temporally precise neural marker of cognitive distraction that closely tracks the latency of multichannel scalp EEG. At the same time, the scalp topographies indicate that the neural signals underlying decoding in both modalities likely capitalize on eye- and head-movement-related processes, highlighting a shared neurobehavioral basis of distraction-related neural signatures.

## Discussion

The present work demonstrates that distraction-related cognitive load during naturalistic driving can be decoded with high fidelity using a single-channel in-ear EEG. Utilizing minimally invasive ear-EEG sensor achieved classification performance and detection latencies on par with a full-cap 24-channel EEG system. The temporal dynamics of the decoding were very similar between the ear electrode and the full scalp montage- both in terms of when distraction-related neural signals emerged and how long they remained informative. Moreover, both systems showed comparable generalization profiles, indicating that the ear-EEG captures essentially the same underlying distraction markers as the conventional EEG. In addition to neural measures, we observed that purely behavioral signals, namely eye movements and head rotations, carried reliable signatures of driver distraction. In particular, eye velocity proved to be a strong early indicator of distraction, with decoding that generalized across time, underscoring the tight link between oculomotor behavior and cognitive load in driving.

The present findings align with and extend prior research using EEG to monitor driver attention and mental workload. Numerous studies have shown that EEG activity -especially in certain frequency bands -is modulated by driver distraction or increased cognitive load(e.g. (Ronca et al., 2024)). Traditional EEG-based approaches to detect distraction have typically employed multi-channel recordings in simulated or controlled driving tasks. These have demonstrated that machine-learning classifiers can distinguish distracted vs. focused states above chance levels using spectral, connectivity, or event-related EEG features (e.g. (Sonnleitner et al., 2014a; Wang et al., 2018; Peng et al., 2022; Hussein et al., 2023; Li et al., 2023; Ronca et al., 2024)). The present use of a single in-ear channel represents an extreme reduction in setup complexity -yet still achieves robust decoding. This supports the notion that even highly pared-down EEG systems can yield meaningful information about driver state. Notably, a recent study by Fan et al. collected EEG from only two forehead electrodes in a train-driving simulator and was able to detect fatigue and distraction, hinting at the viability of low-density EEG for real-world vigilance monitoring (Fan et al., 2022). The present results confirm in a driving context what emerging work in other domains has suggested: ear-EEG signals can reflect cognitive state changes nearly as well as conventional scalp EEG(Crétot-Richert et al., 2023). Ear-centered electrodes could differentiate levels of workload with accuracy comparable to a 12-channel cap, especially when multiple ear electrodes were used (Crétot-Richert et al., 2023). The present study extends this evidence by showing such performance is achievable even with a *single* ear electrode during an ecologically valid driving task.

Is there an additive value of full cap EEG in detecting driver’s distraction? Full EEG caps or at least several electrodes often require substantial preprocessing to manage noise. In particular, a recent systematic review of EEG-based driver state monitoring concluded that, despite extensive methodological development and frequently high reported classification accuracies, existing approaches remain largely ineffective for real-world deployment (Hussein et al., 2023). The primary reasons include reliance on high-density EEG montages, extensive preprocessing pipelines, computationally demanding feature extraction, limited ecological validity, and unclear interpretability of the decoded signals. In this context, the present findings do not contradict the review’s conclusion, but rather offer a complementary view on which aspects of EEG-based decoding may nevertheless be viable in practice. First, the present results directly address the issue of *hardware complexity*. Whereas much of the reviewed literature relies on multi-channel EEG systems without clear justification for electrode selection, our comparison shows that a single in-ear EEG channel yields decoding latencies and temporal generalization profiles that closely match those of a full-cap EEG system. Importantly, the additional value of the scalp montage lies primarily in spatial interpretability rather than in earlier or stronger decoding performance. This distinction is rarely made in prior work and suggests that dense EEG arrays may be more useful for mechanistic interpretation than for real-time detection, a point that is highly relevant for practical applications. Second, the present study responds to concerns regarding *computational burden and preprocessing requirements*. Many approaches reported in earlier work depend on extensive artifact rejection, frequency decomposition, handcrafted features, or deep learning architectures that are difficult to implement in real time. In contrast, the present decoding approach relied on time-resolved signals with minimal preprocessing (i.e. data segmentation, 45 Hz low pass filter and baseline correction) and standard classification methods, yet still achieved reliable discrimination of distraction. This demonstrates that at least part of the relevant information is present in the raw temporal structure of the signals and can be accessed using fast, off-the-shelf pipelines. Third, the multimodal comparison conducted here provides an important reframing of the issue of *ocular and movement-related contributions to EEG decoding*, which are often treated as confounds in the reviewed literature. By explicitly showing that eye velocity and head rotation carry robust and temporally stable distraction-related information, and that the temporal dynamics of ear EEG and scalp EEG closely align with these behavioral measures, the present results suggest that neural decoding in wearable EEG systems likely exploits shared neurobehavioral dynamics associated with visual attention and orienting behavior. Rather than undermining EEG-based approaches, this coupling helps explain why minimal EEG configurations can succeed and why attempts to completely eliminate such signals may be both impractical and counterproductive for real-world monitoring. Finally, the present findings help refine the interpretation of “ineffectiveness” raised in the review literature. While it remains true that many EEG-based driver monitoring systems are currently unsuitable for deployment due to complexity and lack of robustness, the present results indicate that *ineffectiveness is not inherent to EEG as a modality*, but rather to how it has traditionally been implemented and analyzed. When EEG is treated as part of a multimodal, temporally resolved system and when the focus shifts from exhaustive neural isolation to reliable state detection, single-channel ear EEG emerges as a viable compromise between signal fidelity, user acceptability, and computational feasibility. In this sense, the present work complements existing work by identifying a narrower but more practical operating point for EEG-based distraction decoding. It suggests that future progress in the field may come less from increasing model complexity or channel count, and more from embracing minimal, wearable sensors combined with fast, interpretable decoding strategies that align with the behavioral dynamics of distraction itself.

### Multimodal Indicators of Distraction

An important insight from the present results is the utilization of multimodal approaches to distraction detection. Eye movement metrics and head rotations, recorded in parallel with EEG, carried their own reliable signatures of distraction. This is consistent with extensive work highlighting how driver inattention manifests in observable behaviors: for instance, cognitive distraction leads to more rigid visual scanning, fewer mirror checks, and longer glances away from the road (Peng et al., 2025). In the present data, eye velocity in particular provided an early-warning signal of impending distraction- it was among the first measures to indicate a change in the driver’s focus. Prior studies have likewise reported that eye tracking alone can identify driver distraction with reasonably high accuracy (e.g., ∼85 % using gaze dispersion features) in a naturalistic driving context (Musabini and Chetitah, 2020). Head movement cues (such as changes in yaw or roll) add further context, and combining them with eye data yields better performance than either alone. Multimodal integration of physiological and behavioral signals has repeatedly been shown to improve the robustness of driver-state detection. For example, systems combining ear-EEG with cardiovascular measures such as ECG and PPG have achieved very high classification performance for detecting drowsiness, often exceeding 97% accuracy under optimized conditions and long analysis windows, with multimodal fusion yielding more stable performance than any single modality alone(Hong et al., 2018). Importantly, however, such performance gains typically rely on extended temporal windows, modality-specific preprocessing, and feature extraction pipelines that limit real-time applicability. The present work deliberately adopts a different perspective. Rather than maximizing classification accuracy through multimodal fusion, we focus on identifying distraction-related signals that are *early, temporally precise, and accessible using minimal hardware and off-the-shelf decoding methods*. Within this constraint, we show that single-channel in-ear EEG captures distraction-related dynamics with latencies and temporal generalization properties comparable to full-cap EEG, without requiring additional sensors or computational overhead. While multimodal fusion remains a promising direction for future systems, the current results establish a lower-bound, reliable decoding of cognitive distraction using a single wearable neural sensor and fast analysis, consistent with the practical goals outlined in the introduction and discussed below.

### Minimal-Intrusion EEG and Practical Benefits

A key motivation for using in-ear EEG is the practical advantage of minimal intrusion. The presented results demonstrate that a discreet earbud-style EEG sensor can offer a temporally precise neural index of distraction without the cumbersome setup of a full EEG cap. This finding is especially promising in light of recent advancements in wearable EEG technology (Kaongoen et al., 2023; Niso et al., 2023; Sugden et al., 2023; Mathewson et al., 2024). Research has developed around-the-ear electrode arrays and ear canal sensors that record EEG inconspicuously, suitable for everyday environments(Crétot-Richert et al., 2023), that enable to perform ambulatory EEG with electrodes hidden around the ear and a smartphone amplifier, illustrating that high-quality brain signals can be captured on the move(Debener et al., 2012; Debener et al., 2015). The ear region offers several advantages: it is relatively free of hair (improving electrode contact), and a well-fitted earpiece is stable against motion artifacts. By capitalizing on this technology, the present study’s approach remains almost invisible and does not encumber the driver -a stark contrast to traditional EEG hardware. This minimal intrusion is critical for any real-world deployment; a method to monitor distraction is far more likely to be adopted if it can be embedded into something drivers already wear (e.g. a communications earpiece or noise-canceling earbud). Importantly, this is not to suggest that in-ear EEG is inherently superior to other wearable EEG approaches, each of which offers distinct advantages in terms of spatial coverage and signal characterization; rather, from the perspective of readiness for real-world deployment and unobtrusive long-term use, earbud-based EEG represents a particularly practical compromise. Furthermore, the present decoding pipeline was intentionally kept simple, using fast, off-the-shelf machine learning methods with minimal preprocessing. The success of such a lightweight approach implies that real-time implementation is feasible.

### Role of Oculomotor Signals in EEG Decoding

It is worth discussing the contribution of oculomotor processes to EEG-based decoding, as this bears on the interpretation of “what” the ear-EEG is detecting. Driver distraction is inherently linked to eye activity- when a driver’s mind wanders or they engage with a secondary task inevitably introducing ocular artifacts propagating to the simultaneously recorded EEG. In the present case, the in-ear electrode is positioned such that it can pick up some of these eye-related signals in addition to brain activity. Yet, this is not considered a confound or noise to be eliminated; rather, it is a reflection of the coupled neurobehavioral dynamics of distraction. The onset of distraction involves both cortical changes and behavioral output (e.g. gaze shifts), and EEG will inevitably capture a mixture of both. In fact, the ability of EEG -especially from frontal or ear locations -to leverage eye movement signatures is part of its strength in detecting driver state. Prior studies have noted that cognitive distraction often manifests in eye metrics well before it impairs driving performance(Peng et al., 2025). Thus, if the present ear-EEG decoding “capitalizes” on eye movement-related signals, it is effectively incorporating an early-warning aspect of distraction. In the present work, this is not considered as artifactual contamination, but as EEG picking up a valid physiological component of distraction. From a neuroscience perspective, one may wish to disentangle neural activity from peripheral signals; attempts to do so using an overlapping dataset has been reported elsewhere (Popov et al., 2025). For distraction monitoring, however, the primary objective is timely and reliable detection, and in this context the brain’s control of eye movements constitutes an integral component of the distraction phenotype.

In conclusion, the present results demonstrate that an earbud EEG can decode driver distraction with accuracy and temporal resolution comparable to a full EEG cap, while offering the practical advantages of comfort and ease-of-use. Eye movement and head rotation signals further strengthen the detection, and their contribution highlights the integrated nature of cognitive and sensorimotor changes during distraction. While certain limitations and challenges remain -including individual calibration and on-road validation - the evidence presented here supports the ear-EEG as a viable marker of distraction. This low-burden approach could be translated into real-world driver monitoring applications in the near future, helping to bridge the gap between laboratory cognitive neuroscience and everyday safety-critical settings.

## Limitations

Despite the encouraging results, several limitations of the present work should be acknowledged.

### Lack of subject-specific modeling

Subject-specific classifiers were not developed or evaluated in the present study. All decoding analyses were conducted using pooled data with cross-validation across participants, an approach chosen to balance statistical power against the limited number of trials available per individual. Obtaining sufficient within-subject data to support individualized models would require substantially longer driving sessions, which was beyond the scope of the current design but is directly motivated by the present findings. While pooling demonstrates general feasibility, it may obscure individual differences in how cognitive distraction manifests across drivers. Individualized models may yield higher performance by capturing person-specific neural and behavioral patterns, and future studies employing longer recording durations should assess within-subject decoding performance and the benefits of personalization.

### Potential influence of eye-movement-related signals

Part of the discriminative information captured by in-ear EEG likely reflects signals related to eye movements or other peripheral activity. This implies that distraction-related neural activity is intermixed with oculomotor signals, rather than being isolated to purely cortical oscillatory dynamics. Although such contributions are not considered a confound for the purpose of distraction detection, they raise the question of how much of the decoding performance is driven by brain activity versus volume-conducted eye-related signals. Explicit separation or removal of ocular components was not performed in the present analyses. Future work could incorporate independent component analysis or dedicated eye-movement detectors to further characterize the relative contributions of neural and peripheral sources. Nevertheless, for application-oriented monitoring, this blending of signals may be acceptable, as oculomotor control constitutes an integral component of the distraction phenotype.

### Pooled versus individualized classifiers

The use of pooled classifiers trained across multiple participants may limit optimal performance for any given individual, given known inter-individual variability in EEG signals. Cross-subject generalization can be challenging, and models trained on pooled data may fail to capture idiosyncratic features of distraction or, conversely, rely on patterns not expressed uniformly across drivers. Although pooled decoding provides an important proof of feasibility, future implementations may benefit from brief calibration procedures or adaptive learning strategies. Assessing performance on completely unseen individuals and exploring transfer-learning approaches represent important directions for future research.

### Ecological validity and deployment context

The study was conducted in a controlled naturalistic driving environment designed to approximate real-world conditions while maintaining experimental control. Although this setting captures key aspects of driving behavior and distraction, it does not fully reflect the complexity of on-road driving, which includes unpredictable events, variable traffic dynamics, and additional sources of sensory and mechanical noise. This limitation does not undermine the present findings but highlights the need for future validation under fully real-world driving conditions.

## Acknowledgments

This work was supported by the Schweizerischer Nationalfonds zur Förderung der Wissenschaftlichen Forschung (SNF) Grant 105314_207580 awarded to PT and Emma-Louise-Kessler Fund awarded to LS. The authors extend gratitude to all the participants who volunteered for the experiment.

